# Interleukin-16/STAT6 recruits CBP/p300 to upregulate TIMP-3 and promote atherosclerotic plaque stability

**DOI:** 10.1101/2023.05.23.542025

**Authors:** Hui He, Meng Ding, Yuan Zhu, Tianyu Jiang, Doudou Dong, Xiaoting Xu, Hailong Ou

## Abstract

**Background:** Atherosclerotic plaque rupture increases the risk of ischemic heart disease and stroke and commonly causes sudden death. High levels of circulating or intraplaque interleukin 16 (IL-16) are clinically associated with a reduced incidence of cardiovascular events. Here, we investigated the effects of IL-16 on plaque phenotypic modification and identified the molecules involved in smooth muscle cells (SMCs).

**Methods:** We deleted IL-16 in ApoE-/- mice to generate IL16-/-ApoE-/- mice, and the double mutant was used for plaque phenotype analysis after a 24-week high-fat diet. RNA sequencing was performed to identify the changes in cellular processes and molecule expression in response to IL-16 defects. Affinity purification-mass spectrometry was used to identify the STAT6 binding protein. Bone marrow transplantation was used to investigate the effects of hematopoietic IL-16 deficiency or reconstitution on plaque stability.

**Results:** IL-16 deficiency reduced collagen deposition and increased the necrotic core area in the plaques of the brachiocephalic artery and aortic root lesions. Intraplaque TIMP-3 levels were found to be decreased in association with an increase in the proteolytic activity of MMPsinIL16-/-ApoE-/- mice. Next, we demonstrated that IL-16 activates the CD4/JAK2/STAT6 pathway and that STAT6 directly binds the TIMP-3 promoter in SMCs. Furthermore, IL-16 treatment increased the interaction of cAMP-response element binding protein (CBP)/p300 with STAT6, which promoted STAT6 acetylation and increased histone H3 acetylation in the TIMP-3 promoter. Inhibition of CBP/p300 resulted in decreased acetylation of STAT6 and TIMP-3 promoter histone H3 and TIMP-3 expression, suggesting a requirement for CBP/p300 as a coactivator. Finally, we found that hematopoietic-derived IL-16 from ApoE-/- mice or overexpression of TIMP-3 successfully attenuated plaque instability in IL16-/-ApoE-/- mice.

**Conclusions:** IL-6 upregulates TIMP-3 expression and remodels the intraplaque extracellular matrix toward a stable phenotype, suggesting IL-16 as a potential target for intervening in atherosclerosis at later stages.

---

Vulnerable atherosclerotic plaques contain a large necrotic lipid-rich core and high macrophage content with a thin fibrous cap, which is susceptible to rupture, subsequently forming occlusive thrombi and causing acute clinical events such as myocardial infarction and stroke. The fibrous cap is made primarily of SMCs and extracellular matrix (ECM) components, including collagen, elastin, matrix metalloproteinases (MMPs), tissue inhibitors of metalloproteinases (TIMPs) and proteoglycans. In addition to reducing the necrotic lipid-rich core, increasing amounts of ECM collagen and smooth muscle content will stabilize the atherosclerotic plaque and reduce clinical events.

TIMPs are endogenous inhibitors of MMPs that prevent ECM degradation and maintain ECM integrity. TIMP-3, a secreted 24-27-kDa stromal protein, is a member of the TIMP family. Among the four identified mammalian TIMPs (TIMP1-4), TIMP-3 is the only ECM-bound TIMP, whereas TIMP-1 and TIMP-2, as soluble proteins, diffuse freely in the extracellular space. Moreover, TIMP-3 has more broad-spectrum metalloproteinase substrates, which inhibits all MMPs and a number of other metalloproteinases, such as ADAMs (a disintegrin and metalloproteinases) and ADAMTSs (a disintegrin and metalloproteinase with thrombospondin-like motifs).^1,2^ This unique property renders TIMP-3 a major inhibitor of ECM turnover.^1,2^ By interacting with and inhibiting pericellular MMPs, especially MMP2/9, and membrane receptors such as tumor necrosis factor (TNF)-α converting enzyme (TACE)/ADAM17, TIMP-3 profoundly affects vascular remodeling and inflammation. ^2-4^

Interleukin-16 (IL-16) is predominantly produced by immune cells, especially T lymphocytes, and was first described as a lymphocyte chemoattractant factor (LCF). IL-16 is generated by cleaving the precursor molecule pro-IL-16 and functions as a ligand of CD4. By binding to the D4 domain of the CD4 receptor, the bioactive form of secreted IL-16 induces the recruitment, activation and proliferation of CD4-expressing cells, including T lymphocytes, monocytes, eosinophils, dendritic cells, and mast cells, at sites of inflammation during asthma, allergy, and rheumatoid arthritis. ^5-8^ Moreover, IL-16 has been described as a proinflammatory chemokine that amplifies the inflammatory reaction by stimulating the production of multiple cytokines, such as IL-1β, IL-6, IL-15 and tumor necrosis factor (TNF)-α, in activating lymphocytes and other immune cells.^7^ Under certain circumstances, exogenous IL-16 also reduces the production of proinflammatory cytokines, such as in synovial lesions, and antigen-induced airway hyperreactivity, thus exhibiting an immunosuppressive effect.^9-11^

Although the role of IL-16 as a chemoattractant immunomodulatory cytokine in infection and autoimmune diseases has been extensively investigated, the functions of IL-16 in cardiovascular disease are not clear. Elevated IL-16 mRNA in peripheral blood mononuclear cells and increased plasma levels of IL-16 were found in patients with acute myocardial infarction.^12,13^ Cardiac overexpression of IL-16 increased macrophage infiltration and promoted myocardial fibrosis and stiffness.^14^ Neutralizing treatment with an anti-IL16 antibody alleviated cardiac inflammation and fibrosis and improved cardiac dysfunction.^14-16^ This evidence indicates that IL-16 has an adverse effect on cardiovascular function. However, contrary results show that elevated IL-16 in both circulation and carotid plaques is associated with a reduced incidence of cardiovascular events in patients with advanced atherosclerotic lesions.^17,18^ Therefore, the role of IL-16 in cardiovascular diseases remains to be investigated.

In the current study, we emphasize the role of IL-16 in modulating the phenotype of atherosclerotic plaques. Our studies indicate that IL-16 improves intraplaque ECM components and increases plaque stability by upregulating TIMP-3. Mechanistically, we provide evidence that IL-16 activates downstream JAK2/STAT6, which recruits the acetylase CBP/p300 in the nucleus in SMCs. Within the nucleus, STAT6 and CBP/p300 form a complex and bind to the TIMP-3 promoter region, thus initiating TIMP-3 transcription.

## METHODS

Data availability

All supporting data are available from the corresponding author upon reasonable request. A detailed description of the Materials and Methods can be found in the Supplemental Material.

## RESULTS

### IL-16 deficiency reduces atherosclerotic plaque stability

Apolipoprotein E knockout mice (ApoE-/-) and ApoE and IL-16 double knockout mice (IL16-/-ApoE-/-) were fed a high-fat diet (HFD) for 12 and 24 weeks. The body weights and serum lipids, including triglycerides (TGs), total cholesterol (TC), low-density lipoprotein (LDL) and high-density lipoprotein (HDL), were not significantly altered in IL16-/-ApoE-/- mice after 12 and 24 weeks of HFD feeding (Table S1). The plaque burden on the whole aorta and the size of atherosclerotic lesions at the aortic root did not differ between IL16-/-ApoE-/- and ApoE-/- mice fed an HFD for 12 weeks, while aortic root lesions were increased in IL16-/-ApoE-/- mice compared with ApoE-/- mice when the HFD was extended to 24 weeks (Figure S1A, B).

To analyze atherosclerotic plaque stability, we focused on advanced lesions from the brachiocephalic artery (BCA) of mice with 24 weeks of HFD induction. IL-16 deletion resulted in increased lipid accumulation and macrophage infiltration within the plaques (Figure 1A). Intraplaque collagen and SMC contents were dramatically reduced in IL16-/-ApoE-/- mice compared with ApoE-/- mice, thus increasing the plaque vulnerability index (Figure 1A). Moreover, lesions from IL-16-null mice exhibited a larger necrotic core size and thin fibrous cap, two features of plaque instability (Figure 1B). Buried fibrous caps, discontinuous fibrous cap and intraplaque hemorrhage all reflect the occurrence of plaque rupture and were more frequently found in the plaques from IL16-/-ApoE-/- mice relative to ApoE-/- mice (Figure S2). In addition, the plaque phenotype at lesion sites of aortic roots from the two kinds of mice was investigated. A larger necrotic core associated with a thinner fibrous cap and less collagen content was detected within the plaques of IL-16-/-ApoE-/- mice compared with those of ApoE-/- mice after 24 weeks of HFD feeding (Figure 1C). Together, these data revealed that IL-16 deficiency changes atherosclerotic plaque composition toward an unstable phenotype in mice.

**Figure 1.**
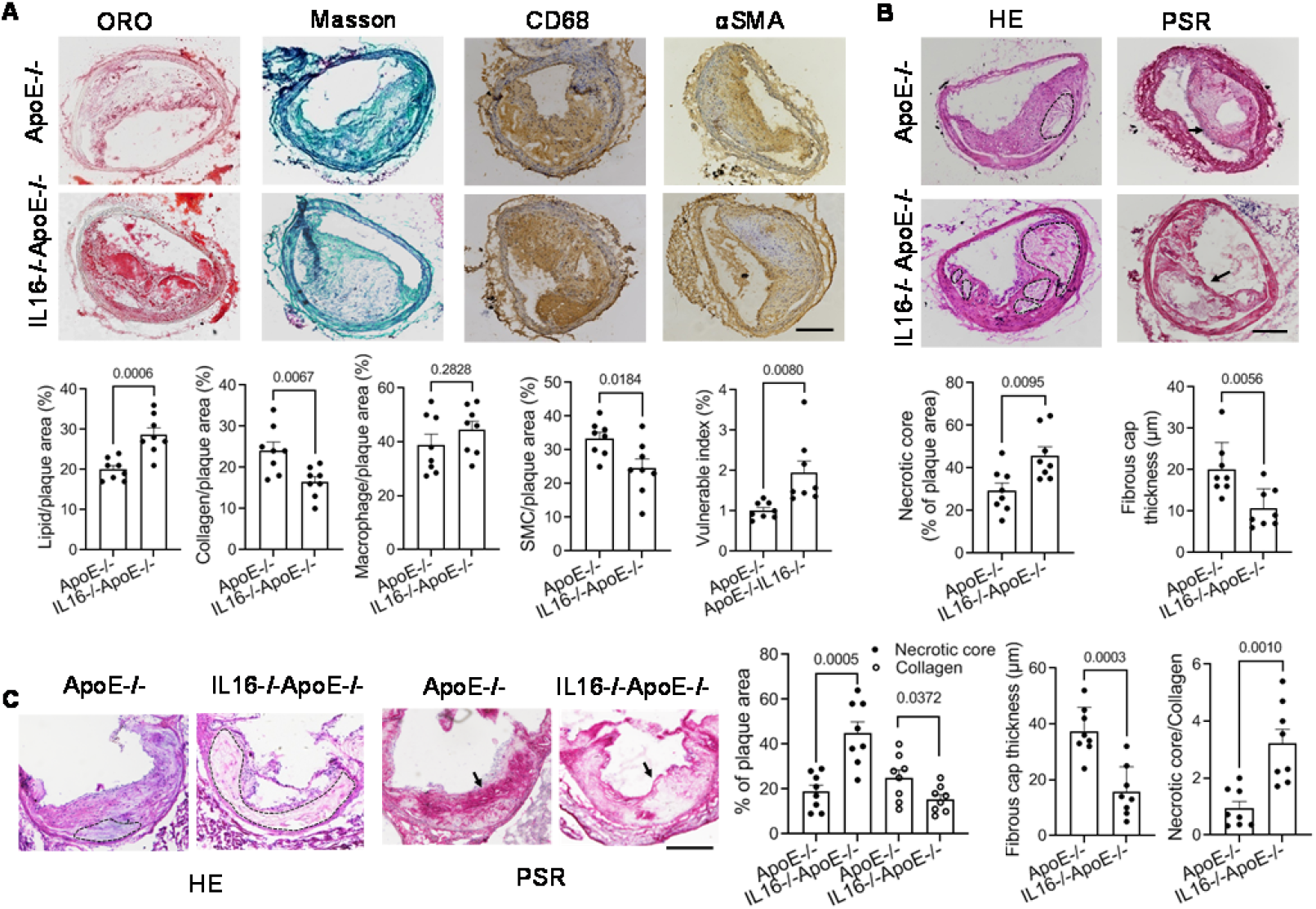
Increased atherosclerotic plaque vulnerability in IL16-/-ApoE-/- mice. IL16-/-ApoE-/- and ApoE-/- mice were fed a high-fat diet at eight weeks of age for 24 weeks. **A**. Frozen sections from brachiocephalic artery (BCA) lesions were stained with oil red O, Masson’s trichrome, anti-CD68 and anti-αSMA antibodies, which were used to indicate lipids, collagen, macrophages and smooth muscle cells, respectively. The staining was quantified by calculating the positive area in the total plaque area. The plaque vulnerability index was determined by (macrophage area + lipid area)/(collagen area + SMC area). **B**. BCA sections were stained with hematoxylin and eosin (HE) and picrosirius red (PSR). Regions resistant to HE staining were defined as necrotic cores and are outlined with black dashed lines. Fibrous cap thickness is indicated by the arrow in PSR-stained sections. **C**. Sections from aortic roots were stained with HE or PSR. The percentage of necrotic core and collagen area in the total plaque area were calculated, and fibrous cap thickness was measured. The data are presented as the mean±SEM, and the *P* value was analyzed by unpaired t test. n=8 mice for each group. Scale bar, 200 μm.

### 2. IL16 deletion promotes intraplaque extracellular matrix degradation

To identify the molecules and pathways involved in IL-16-induced plaque phenotypic modulation, we performed transcriptome screening by poly(A)+ RNA sequencing in aortas from IL16-/-ApoE-/- and ApoE-/- mice after 24 weeks of HFD feeding. Global analysis showed that the expression of a total of 1201 genes was altered, of which 943 genes were upregulated and 258 genes were downregulated in mice with IL-16 deletion (Figure S3A). We performed Gene Ontology (GO) analysis to characterize and functionally cluster these genes and found that genes encoding membrane and extracellular components were remarkably enriched (Figure S3B). Gene set enrichment analysis (GSEA) further revealed downregulated gene signatures of extracellular matrix organization and remodeling in IL-16-deficient mice (Figure 2A; Figure S3C). The heatmap showed that loss of IL-16 broadly reduced the expression of genes related to the ECM, including Collagen I α1 (ColIα1), ColIα2, Elastin, Filamin-α (Flnα), integrin beta 1 (Itgb1) and TIMP-3 (Figure 2B). The aortic expression of ColIα1 and TIMP-3 was further validated by quantitative reverse transcription PCR (qRT‒PCR) and Western blotting and was found to be dramatically reduced at both the mRNA and protein levels in IL-16-null mice (Figure 2C,D). Consistently, there were fewer SMCs expressing TIMP-3 indicated by costaining of TIMP-3 (green) and SMCs (red) in IL16-/-ApoE-/- mice than ApoE-/- mice (Figure 2E). A 42% decrease in the in situ expression of TIMP-3 within the plaques of aortic roots from IL16-/-ApoE-/- mice was detected compared with those of ApoE-/- mice after 24 weeks of HFD feeding (Figure 2E). The intraplaque TIMP-3 level was negatively correlated with the plaque vulnerability in IL16-/-ApoE-/- mice (Figure 2F).

**Figure 2.**
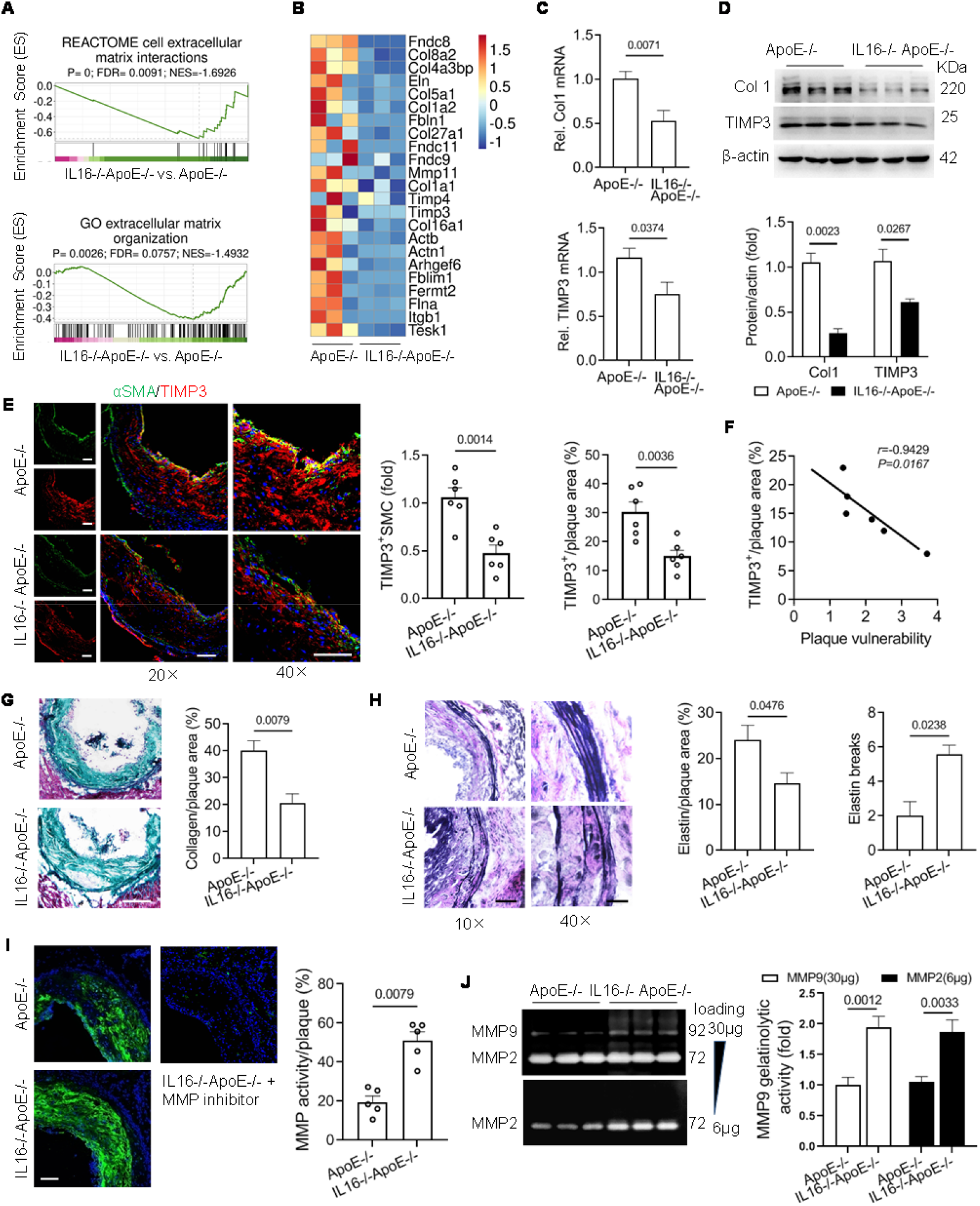
IL-16 deficiency increases MMP activity and extracellular matrix disruption within atherosclerotic plaques. IL16-/-ApoE-/- and ApoE-/- mice were fed a high-fat diet at eight weeks of age for 24 weeks. **A**. Gene set enrichment analysis (GESA) of the differentially expressed genes in the aortic RNA sequencing data. **B**. Heatmap showing the expression of genes encoding extracellular matrix components in the aorta. **C, D**. The mRNA and protein levels of ColIα1and TIMP-3 in the aorta were measured by qRT‒PCR and Western blotting, respectively. n=6 mice/group. Unpaired t test. **E**. Immunofluorescence staining of TIMP-3 (green) and α-SMA (red) in cryosections of aortic roots. The colocalization was indicated as yellow staining. Scale bar, 100 μm. n=6 mice/group. Unpaired t test. **F**. TIMP-3 expression correlated with fibrous cap thickness. n=6 mice/group. Spearman correlation test. **G, H**. Masson’s trichrome and VVG staining of aortic root sections. Scale bar, 200 μm in G, 100 μm in H (left) and 25 μm in H (right). n=5. Mann‒Whitney U test. **I**. MMP proteolytic activity in aortic root plaque sections was measured by in situ zymography using fluorescein-conjugated gelatin as a substrate. Ilomastat was used to inhibit enzymatic activity. Scale bar, 100 μm. n=5 mice/group. Mann‒Whitney U test. **J**. Gelatin zymography analysis of MMP activity in isolated aortas. The band intensities indicated the activity of MMP9 (upper panel, loading amount 30 μg) and MMP2 (bottom panel, loading amount 6μg). n=6 mice/group, unpaired t test.

Collagen and elastin fibers are two major ECM components in vessels and are key determinants for plaque stability. As revealed in Figure 1D, the intraplaque collagen content assessed by picrosirius red staining was 33% lower in IL16-/-ApoE-/- mice than in ApoE-/- mice. Masson’s trichrome staining further confirmed the lower collagen accumulation within the atherosclerotic lesions in IL16-/-ApoE-/- mice (Figure 2G). Elastin was reduced by 41% in IL16-/-ApoE-/- mice compared to ApoE-/- mice and showed irregular arrangement with an increased number of breaks in the vascular wall in IL-16-deficient mice (Figure 2H). Consistently, the results from in situ zymography showed that the defect in IL-16 caused a 1.5-fold increase in the proteolytic activity of MMPs in aortic root lesions, as evidenced by green fluorescence intensity (Figure 2I). Gelatin zymography further showed stronger MMP2 and MMP9 gelatinolytic activities in aortas isolated from IL16-/-ApoE-/- mice than in aortas isolated from ApoE-/- mice (Figure 2J). Collectively, the data indicated that loss of IL-16 reduced the expression of genes involved in ECM remodeling and promoted ECM degradation.

### 3. IL-16 increases JAK2/STAT6-mediated TIMP-3 expression in VSMCs

The Janus kinase (JAK)/signal transducers and activators of transcription (STAT) intracellular signaling pathway is commonly activated in response to cytokine stimulation. As expected, we found that IL-16 generally increased JAK1/2/3 and STAT1/3/5/6 phosphorylation in the rat smooth muscle cell line A7R5 in the time course analysis (Figure S4A, B). Among them, JAK2 and STAT6 showed stronger phosphorylation by IL-16 treatment, and dose-dependent increases at concentrations of 0-100ng/mL (Figure S4C). The increased phosphorylated JAK2 (p-JAK2) and phosphorylated STAT6 (p-STAT6) induced by IL-16 were confirmed in primary mouse VSMCs (Figure 3A, B). Furthermore, we isolated aortas and found that p-JAK2 and p-STAT6 were dramatically reduced in IL16-/-ApoE-/- mice compared with ApoE-/- mice (Figure 3C). The levels of p-JAK2 and p-STAT6 within aortic plaques evaluated by immunohistochemical analysis showed significant reductions in the sections from IL-16 knockout mice (Figure 3D). Thus, the in vitro and in vivo data indicated that JAK2/STAT6 was involved in the IL-16 signaling pathway in smooth muscle cells.

**Figure 3.**
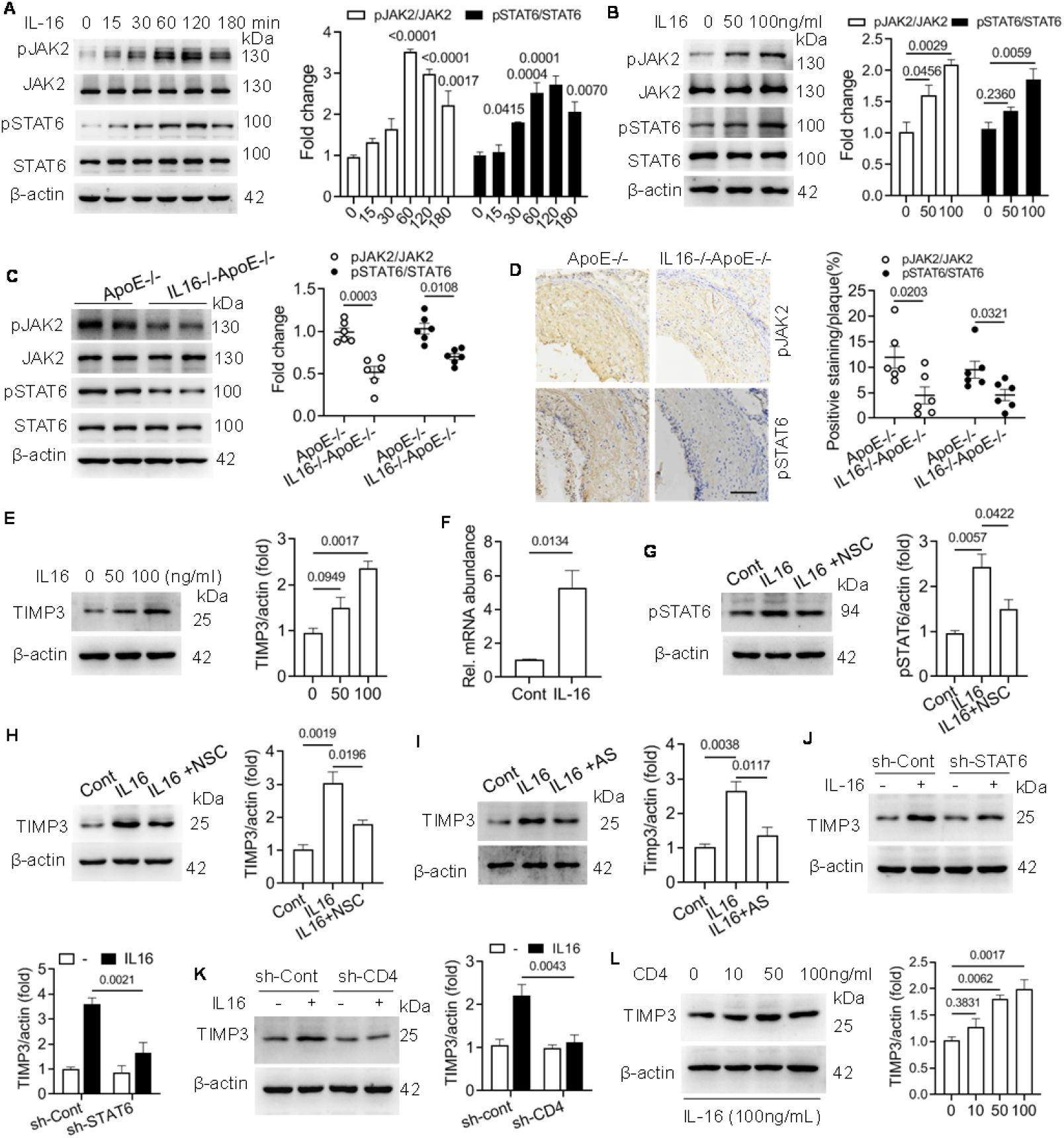
IL-16 stimulates the JAK2/STAT6 pathway and upregulates TIMP3. **A**. VSMCs isolated from wild-type mice (WT) were treated with IL-16 (100 ng/mL) for the indicated minutes. Phosphorylated JAK2 (pJAK2) and STAT6 (pSTAT6) were detected by Western blotting. Values were compared with the control (0 minutes). **B**. Levels of pJAK2 and pSTAT6 measured in isolated VSMCs incubated with 50 or 100 ng/mL IL-16 for 24 h. **C**,**D**. pJAK2 and pSTAT6 in aortas from IL16-/-ApoE-/- and ApoE-/- mice were detected by immunoblotting and immunohistochemical staining. Scale bar, 100 μm. n=6 mice/group, unpaired t test. **E, F**. Protein and mRNA levels of TIMP-3 in VSMCs incubated with IL-16 (50 and 100 ng/mL) for 24 h. n=3, t test in F. **G, H**. Western blotting detection of pSTAT6 and TIMP-3 expression in VSMCs incubated with the JAK2 inhibitor NSC42834. **I**. TIMP-3 expression in VSMCs incubated with the STAT6 inhibitor AS1517499. **J**,**K**. TIMP-3 expression in VSMCs with genetic knockdown of STAT6 (shSTAT6) or CD4 (shCD4). n=3, two-way ANOVA. **L**. TIMP-3 expression in VSMCs with recombinant CD4 knockdown at various concentrations. All the data were obtained from three independent repeats (n=3) and analyzed withone-way ANOVA unless otherwise indicated in C, D, F, J and K.

Having established that JAK2/STAT6 and the ECM remodeling genes TIMP-3 and collagen I were promoted by IL-16, we then explored whether JAK2/STAT6 directly upregulated these genes. Given that the transcriptional regulation of TIMP-3 has been relatively poorly characterized compared to that of type I collagen and that TIMP-3 plays a predominant role in maintaining ECM homeostasis among the TIMPs, we will focus on TIMP-3 and examine its expression controlled by IL-16 in more detail in the following study. First, mouse VSMCs were treated with IL-16 (50-100 ng/mL), and TIMP-3 at both the mRNA and protein levels was found to be dose-dependently increased (Figure 3E,F). Similar results were obtained in human aortic smooth muscle cells (HASMCs) and the A7R5 cell line (Figure S4D, E). Next, we blocked JAK2 with the pharmacological inhibitor NSC42834 and found that both p-STAT6 and TIMP-3 were reduced in VSMCs (Figure 3G,H). TIMP-3 expression was also decreased in VSMCs upon treatment with the STAT6 inhibitor AS1517499 and in cells with genetic knockdown of STAT6 (Figure 3I,J). Finally, we found that reducing membrane receptor CD4 by RNA knockdown attenuated TIMP-3 expression in IL-16-treated VSMCs; supplementation with exogenous CD4 promoted its expression (Figure 3K,L). Together, these results demonstrated that TIMP-3 expression induced by IL-16 is dependent on the receptor CD4 and the downstream JAK2/STAT6 pathway.

### 4. STAT6 directly upregulates TIMP-3 transcription in IL16-induced VSMCs

To analyze TIMP-3 transcription controlled by IL-16/STAT6, we generated a luciferase expression vector containing the -1.9 kb TIMP-3 promoter and transfected it into VSMCs. Reporter assays showed that transcriptional activity was stably increased with increasing concentrations of IL-16 (50-100 ng/mL) (Figure 4A). This response was sensitive to STAT6 inhibition by AS1517499 (Figure 4A). To map the regulatory elements, we generated luciferase reporters with a series of 5’-truncated TIMP-3 promoters (Figure 4B, left). The vectors were transfected into SMCs with or without IL-16 stimulation. The results from the luciferase assay revealed that IL-16 could only significantly transactivate the promoter that was longer than 800 bp upstream of the transcription start site, which harbored the STAT6 consensus recognition sequence located at -823 nt and -1676 nt (Figure 4B, right; Figure S5A). Site-directed mutagenesis of the two sites dramatically abolished IL-16-induced TIMP-3 transcriptional activity, as indicated by luciferase activity assays (Figure 4C). To provide direct evidence of STAT6 binding to the TIMP-3 promoter, we performed EMSAs using a nucleotide probe containing STAT6-binding elements at -823 nt in the TIMP-3 promoter (Figure S5A). The interaction between recombinant STAT6 protein and the probe was apparently detected in VSMCs (Figure 4D). An EMSA assay was further performed in SMCs treated with 50 and 100 ng/mL of IL-16, and the IL-16-induced endogenous STAT6 at both doses in nuclear extracts was shown to specifically bind the nucleotide probe (Figure 4E, left). The presence of STAT6 in the binding complex was confirmed by the addition of STAT6 antibody that caused a supershift (Figure 4E, right). Finally, we confirmed the in vivo TIMP-3 promoter DNA/STAT6 binding by chromatin immunoprecipitation (ChIP) assays, in which IL-16 at 100 ng/mL caused a 3.2-fold increase in the binding activity at -823 nt (Figure 4F,G). We also found that IL-16 increased the binding activity of STAT6 with another STAT6-binding element at -1676 nt in the TIMP-3 promoter in the EMSA but not in the ChIP assay (Figure S5A-D). Together, the data demonstrated that STAT6 promoted TIMP-3 transcription by directly binding to its promoter.

**Figure 4.**
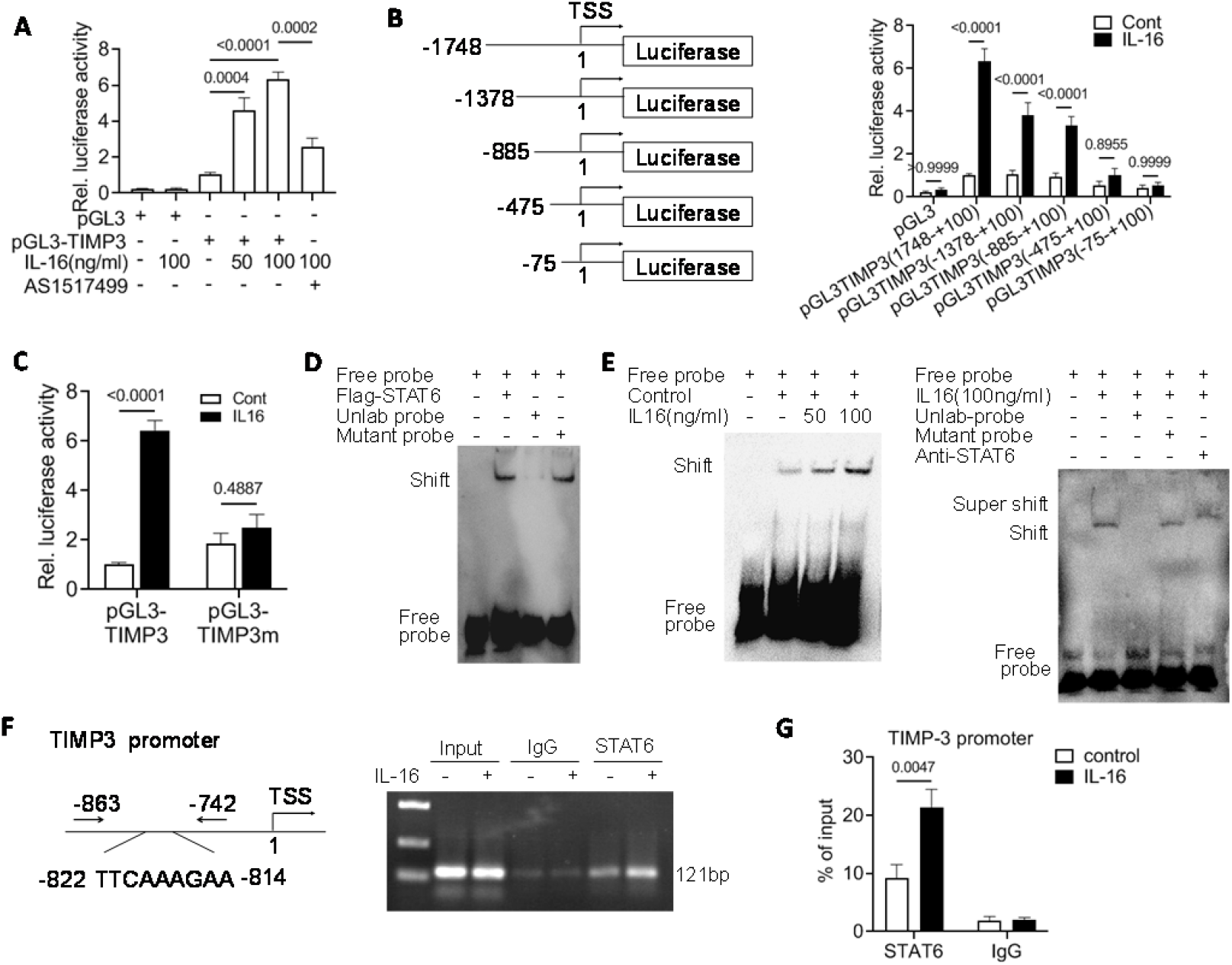
IL-16 increases STAT6 binding to the TIMP3 promoter and enhances promoter activity. **A**. Luciferase reporter assay of TIMP3 promoter activity. VSMCs were transfected with the blank plasmid pGL3 or pGL3-TIMP3 containing the -1.9 kb TIMP-3 promoter sequence and incubated with IL-16 (0, 50, 100 ng/mL) and AS1517499. n=3, one-way ANOVA with Tukey’s pairwise test. **B**. A series of 5’-deletion mutants of TIMP-3 promoters (left) were transfected into VSMCs, and their activities in response to IL-16 stimulation were determined by luciferase reporter assays of VSMCs. n=3, two-way ANOVA with Sidak’s test. **C**. The luciferase activities of the TIMP-3 promoter with mutation of two STAT6 consensus recognition sites (−823 bp and -1676 bp). n=3, two-way ANOVA with Sidak’s test. **D**. EMSA for the binding of recombinant STAT6 to the TIMP-3 promoter. A biotin-labeled oligonucleotide containing the STAT6 binding element at -823 nt in the TIMP-3 promoter was used as a probe. **E**. The binding of the nuclear extract of IL-16 (0, 50, 100 ng/mL)-treated VSMCs to the probes at -823 nt of the TIMP-3 promoter. The presence of STAT6 was detected by incubation with STAT6 antibody and was indicated in the supershift band, and the specificity of probe binding was verified by adding unlabeled or mutant probes. **F, G**. ChIP analysis of STAT6 binding to the TIMP-3 promoter in VSMCs treated with or without IL-16. The primers designed are schematically illustrated in the left panel. The results were detected by agarose gel electrophoresis and quantitative PCR (qPCR). n=3,two-way ANOVA with Sidak’s test.

### 5. IL-16 recruits CBP/p300 and acetylates STAT6 in VSMCs

To gain more insight into the mechanism controlling the STAT6-mediated TIMP-3 transcript, we stably transfected VSMCs with Flag-tagged STAT6, followed by IL-16 stimulation. The STAT6-containing protein complex was affinity purified from the cell extracts with the anti-FLAG antibody and visualized by silver staining (Figure 5A). Mass spectrometry analysis identified cAMP-response element binding protein (CBP) and its homolog p300, a known protein lysine acetyltransferase, in the complex. Western blotting of the column-bound proteins with anti-CBP antibody also confirmed the presence of CBP/p300 in the STAT6 complex (Figure 5B). We performed coimmunoprecipitation to assess the in vivo association of STAT6 and CBP/p300 in IL-16-stimulated cells. The results showed that Flag-STAT6 could be coprecipitated with HA-CBP in transfected cells, and the interaction was strikingly increased after IL-16 treatment (Figure 5C). IL-16 was also demonstrated to promote the binding of endogenous STAT6 and CBP by reciprocal coimmunoprecipitation assays (Figure 5D). Furthermore, recombinant glutathione S-transferase (GST)-fused STAT6 protein expressed and purified from bacteria successfully pulled down CBP in the extract of IL-16-exposed cells, suggesting the in vitro interaction of STAT6 and CBP (Figure 5E). We purified a series of deletion mutants of GST-fused CBP proteins from bacteria and incubated them with cell extract after IL-16 treatment (Figure 5F). The results from the pulldown assay showed that the C/H3 domain (Lane 5) was required for CBP binding with STAT6 (Figure 5F). Finally, Flag-STAT6-transfected VSMCs were stimulated with IL-16, and cell lysates were immunoprecipitated with an anti-Flag antibody. The results from immunoblotting with an antibody against acetyl-lysine showed that IL-16 significantly induced the acetylation level of STAT6 (Figure 5G). Similarly, we immunoprecipitated with an anti-STAT6 antibody in cells treated with or without IL-16 and found IL-16 increase endogenous STAT6 acetylation (Figure S6). Genetic knockdown of CBP abolished IL-16-induced STAT6 acetylation as well as TIMP-3 expression (Figure 5H,I). Consistently, STAT6 acetylation and TIMP-3 expression were reduced when the cells were treated with C646, a pharmacological inhibitor of p300 (Figure S7A,B). Thus, we identified CBP/p300 as a coactivator recruited in the nucleus that promotes STAT6 acetylation, which leads to TIMP-3 upregulation.

**Figure 5.**
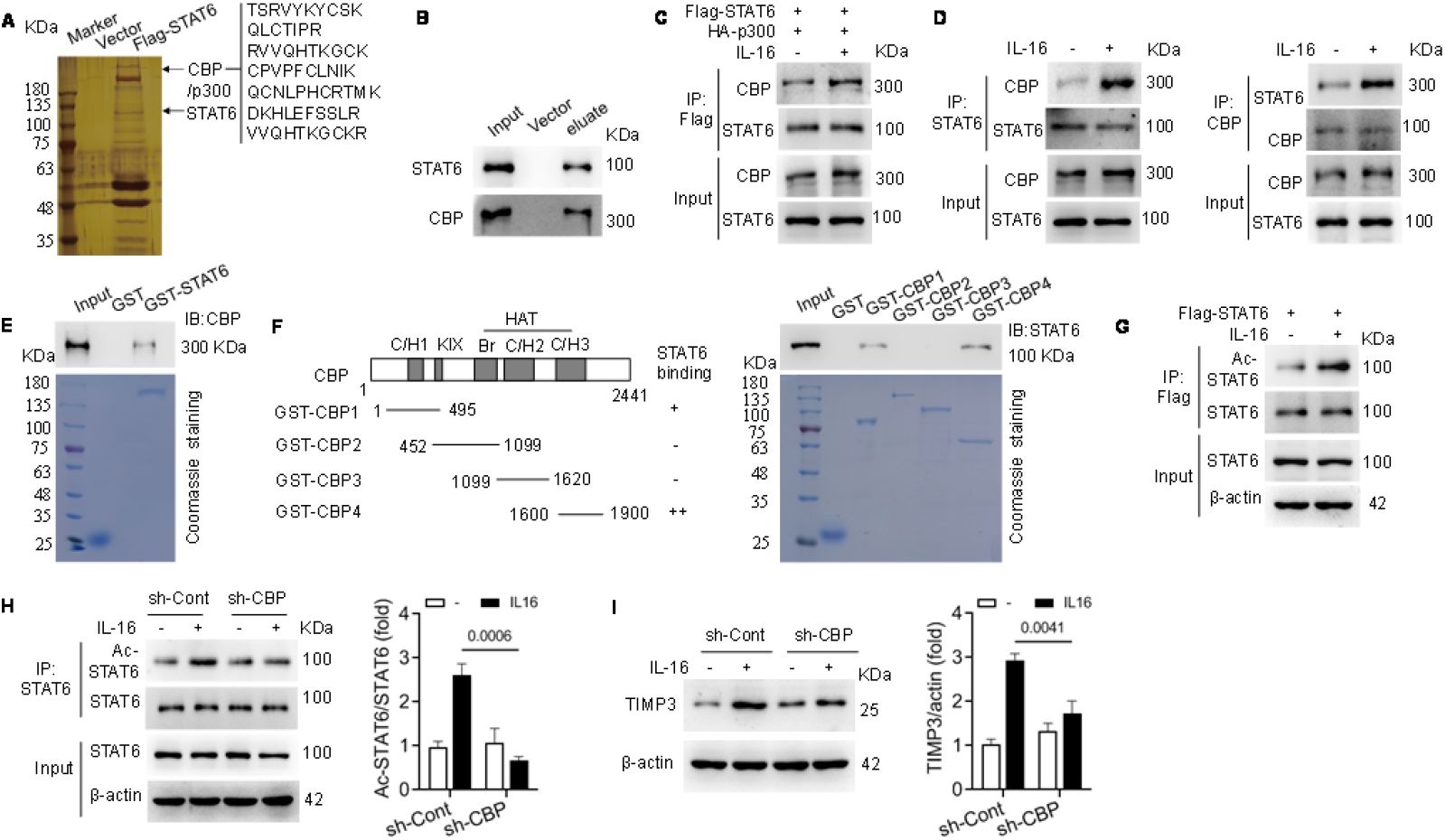
IL-16 increases STAT6 acetylation by recruiting CBP/p300 in VSMCs. **A**.Analysis of STAT6-associated protein.VSMCs were transfected with Flag-STAT6 and treated with IL-16 for 24 h. After affinity purification with anti-FLAG affinity columns, the eluates containing Flag-STAT6 were resolved by SDS‒PAGE and silver stained. STAT6-associated protein was analyzed by mass spectrometry. **B**. Column-bound protein was immunoblotted with CBP antibody. **C**. VSMCs were cotransfected with Flag-STAT6 and HA-CBP following IL-16 incubation. The interaction of exogenous STAT6 and CBP was analyzed by coimmunoprecipitation assays. **D**. Reciprocal coimmunoprecipitation assays for the binding of endogenous STAT6 and CBP in IL-16-induced VSMCs. **E**. GST pulldown assay for the in vitro interaction of STAT6 and p300. Bacterially purified GST-STAT6 was incubated with cell lysates from IL-16-treated cells, followed by GST pulldown and immunoblotting with anti-CBP. The purified GST and GST-STAT6 proteins were detected by Coomassie blue staining. **F**. Mapping the interaction domain of CBP with STAT6. Schematic illustration of various CBP deletion mutants. GST and GST-fused CBP mutants were stained with Coomassie blue. The binding protein was immunoblotted with anti-STAT6. **G**. Isolated VSMCs were transfected with Flag-STAT6 and incubated with or without IL-16, and cell lysates were immunoprecipitated with a anti-Flag antibody. STAT6 acetylation levels were determined by immunoblotting analysis. **H**,**I**. Immunoblotting analysis of STAT6 acetylation and TIMP-3 expression in CBP knockdown VSMCs stimulated with IL-16. n=3, two-way ANOVA with Sidak’s test.

### 6. IL-16 increases CBP/p300-mediated acetylation of TIMP-3 promoter histones in VSMCs

Given that CBP/p300, as a known acetyltransferase, catalyzes protein and DNA acetylation, we then question whether the recruitment of CBP/p300 alters histone acetylation levels in the TIMP-3 promoter. The results from the ChIP assay showed that IL-16 treatment significantly increased CBP/p300 enrichment in the TIMP-3 promoter regions compared with that of the untreated VSMCs (Figure 6A). The acetylation at histone H3 Lys9, 18 and 27 in the TIMP-3 promoter was higher after IL-16 incubation (Figure 6B). Furthermore, the increase in H3 histone acetylation after IL-16 treatment was attenuated when CBP/p300 was knocked down (Figure 6C). Similarly, C646 treatment also reduced H3 histone acetylation levels (Figure 6D). We disrupted the JAK2/STAT6 pathway with the JAK2 inhibitor NSC42834 in IL-16-treated cells and found that the acetylation of STAT6 and TIMP-3 promoter H3 was dramatically decreased (Figure 6E,F). We further found that the association of p300 with STAT6 was increased in the nucleus, as expected, but this did not occur in the cytoplasm (Figure 6G,H). Therefore, these data indicate that CBP/p300 is recruited by STAT6 and forms a complex in the nucleus after IL-16 treatment, while the interaction with CBP/p300 in turn facilitates STAT6 activation (Figure 6I).

**Figure 6.**
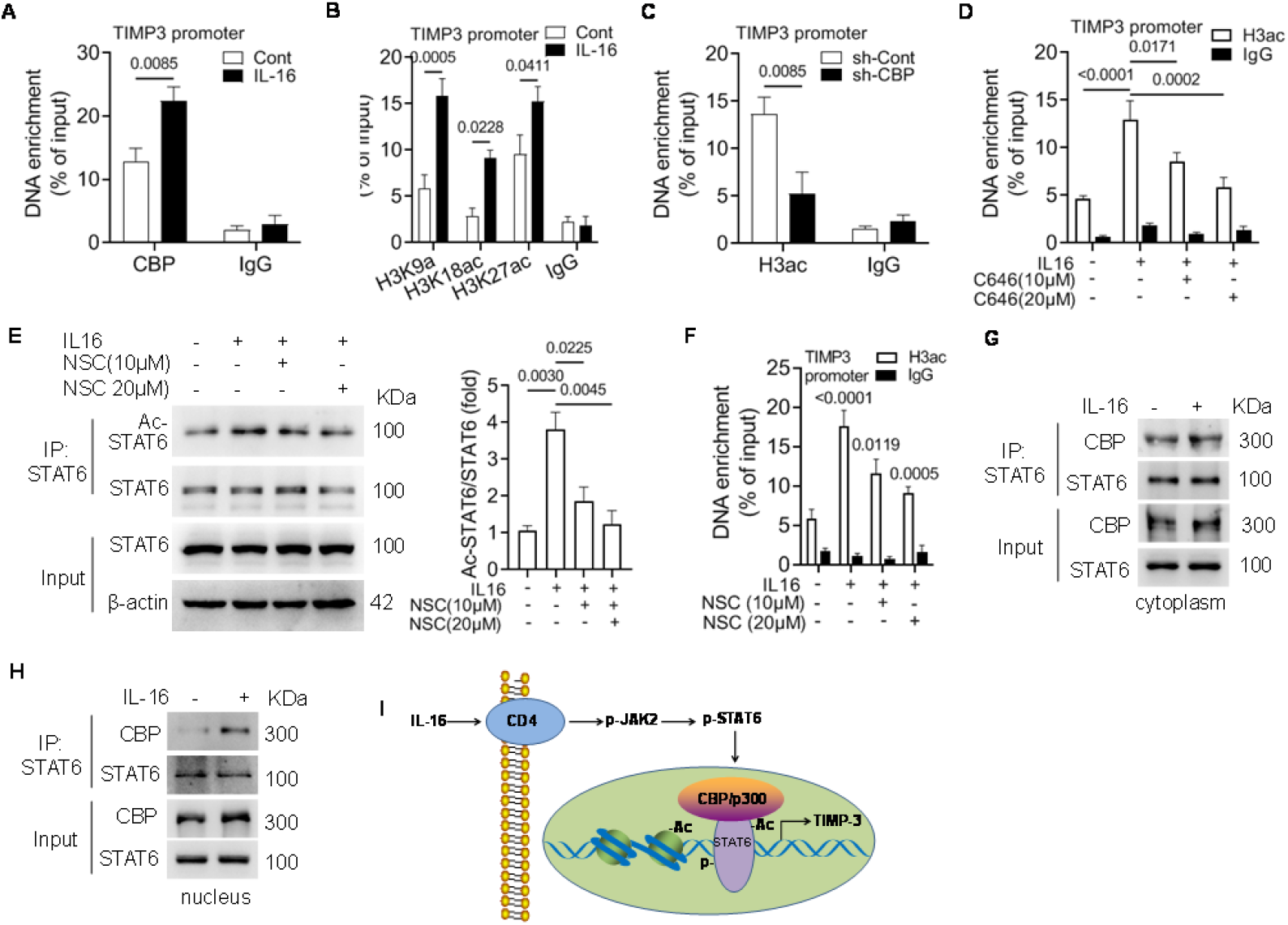
Increased acetylation of TIMP-3 promoter histone by CBP/p300 in IL-16-induced VSMCs. **A**. ChIP assay for the binding of CBP to the TIMP-3 promoter. Cells were incubated with IL-16, and the lysates were precipitated with a CBP antibody. Primers were specific for -822 to -814 bp of the STAT6 binding site. **B**. ChIP assay for histone acetylation with H3K9a, H3K18a, and H3K27a antibodies at the STAT6 binding site in IL-16-induced VSMCs. **C, D**. The level of histone acetylation was assayed by ChIP in IL-16-induced VSMCs with CBP/p300 inhibition by C646 or genetic knockdown. **E, F**. STAT6 and histone acetylation assayed by immunoblotting and ChIP assays, respectively, in IL-16-induced VSMCs treated with or without the JAK2 inhibitor NSC42834. *P*<0.001, compared with the untreated control; *P*=0.0119 and *P*=0.0005, compared with IL-16-treated cells. **G**,**H**. The association of STAT6 and CBP in the cytoplasm and nucleus of IL-16-induced VSMCs. **I**. Schematic representation of pathways for upregulation of TIMP-3 by IL-16. Ac: acetylation; P: phosphorylation. All data are from three independent repeats, two-way ANOVA with Sidak’s test in A-D and F, one-way ANOVA with Tukey’s test in E.

### 7. Effects of hematopoietic IL-16 on plaque stability

We performed bone marrow transplantation (BMT) to further determine the effects of hematopoietic IL-16 deficiency or replenishment on plaque stability. First, we transplanted ApoE-/- or IL16-/-ApoE-/- bone marrow into lethally irradiated recipient ApoE-/- mice (Figure 7A) and found that the mice reconstituted with IL16-/-ApoE-/- BMCs did not have plasma lipid alterations but showed a significant reduction in fibrous cap and collagen deposition and an increase in necrotic core and plaque vulnerability in BCA compared to the mice transplanted with ApoE-/- BMCs (Figure 7B-F; Figure S8, Table S2). Hematopoietic IL-16 deficiency also caused TIMP-3 reduction and an increase in the in situ MMP proteolytic activity within the aortic lesions, in line with the findings in the mouse model of whole-body loss of IL-16 (Figure 7G-I).

**Figure 7.**
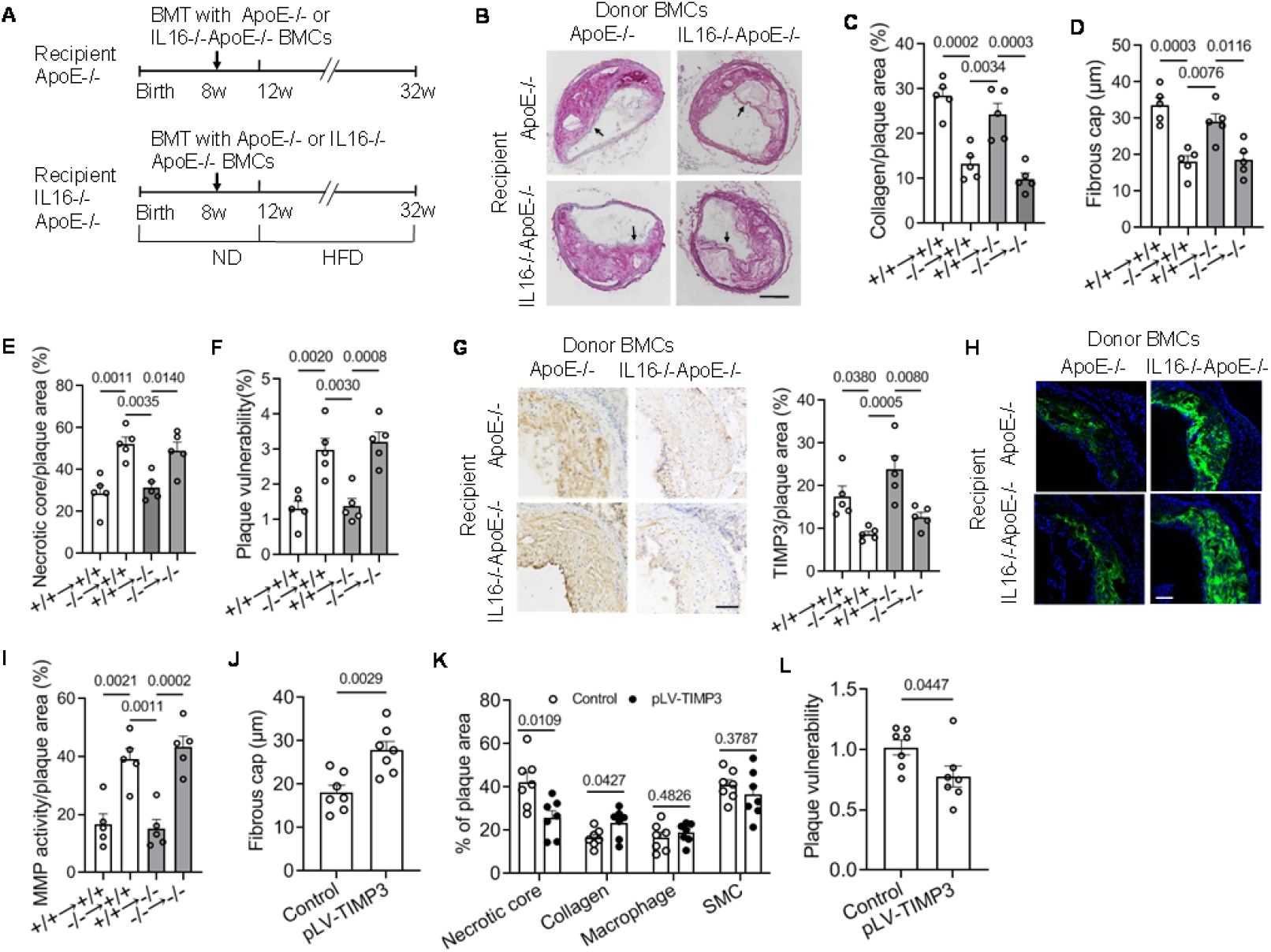
Transplantation with ApoE-/- mouse bone marrow or overexpression of TIMP3 improved the plaque phenotype in IL16-/-ApoE-/- mice. **A**. Strategy for bone marrow cross-transplantation between ApoE-/- and IL16-/-ApoE-/- mice. **B-E**. BCA sections from the four mice created by BMT were stained with picrosirius red and counterstained with hematoxylin. Arrows indicate the fibrous cap. Collagen contents were measured as the percentage of positive PSP area to total plaque area. Necrotic cores of hematoxylin resistant area were quantified. Scale bar, 200 μm. **F**. Plaque vulnerability index of the four BMT mice. **G-I**. TIMP-3 expression and MMP enzymatic activity in the aortic roots of BMT mice measured by IHC staining and in situ zymography. Scale bar, 100 μm. **J-L**. IL16-/-ApoE-/- mice were transfected with lentivirus-mediated TIMP-3 expressing vector (pLV-control) or blank control vector (pLV-control) and subjected to an HFD for 12 weeks. Quantification of the fibrous cap, necrotic core, collagen contents, macrophage and SMC abundance, and plaque vulnerability in the mice transfected with pLV-TIMP3 or control vector. The plaque vulnerability index was determined as percent of the macrophage and necrotic core area in the collagen plus SMC area. In B-I, n=5 mice/group, one-way ANOVA with Tukey’s test; in J-L, n=7 mice/group, unpaired t test.

Next, donor ApoE-/- and IL16-/-ApoE-/- BMCs were transplanted into IL16-/-ApoE-/- mice (Figure 7A). In comparison with that of IL16-/-ApoE-/- BMCs, transfer of ApoE-/- BMCs reduced plaque vulnerability, manifesting enhanced fibrous cap and collagen deposition and a reduced necrotic core, which was associated with increasing intraplaque TIMP-3 and reducing in situ MMP activity without changes in the serum lipid profile (Figure 7B-I; Figure S8, Table S2). The results suggested that hematopoietic reconstitution of IL-16 attenuated plaque instability in IL16-/-ApoE-/- mice.

In addition, there were comparable levels of plaque vulnerability between IL16-/-ApoE-/- and ApoE-/- mice both receiving ApoE-/- BMCs, as well as between IL16-/-ApoE-/- and ApoE-/- mice both receiving IL16-/-ApoE-/- BMCs (Figure 7F). This finding implied that macrophage-derived IL16 mainly contributed to its atheroprotective function.

### 8. Overexpression of TIMP-3 attenuates plaque instability in IL16-/-ApoE-/- mice

We introduced a recombinant lentiviral vector that overexpressed TIMP-3 (pLV-TIMP3) into IL16-/-ApoE-/- mice. After a 12-week high-fat diet, the TIMP-3 level was increased, and the in situ MMP activity was reduced in the plaque of BCA (Figure S9A,B). Overexpression of TIMP-3 caused a more stable plaque phenotype, in which reduced necrotic core and increased collagen content and fibrous cap thickness at BCA were identified in the pLV-TIMP3-infected mice compared with the mice with control vector lacking the TIMP-3 coding sequence (pLV-control) (Figure 7J,K; Figure S9C). The abundance of SMCs and macrophages showed no significant difference within the plaques (Figure S9D). Correspondingly, plaque vulnerability was reduced in the TIMP-3-overexpressing mice (Figure 7L). Therefore, the data confirmed that the beneficial role of IL-16 in atherosclerotic plaque stabilization was dependent on TIMP-3.

## DISCUSSION

Here, we provide compelling evidence that IL-16 depletion causes rearrangement of extracellular matrix components and increases atherosclerotic plaque vulnerability. Furthermore, we found that IL-16 activates the JAK2/STAT6 signaling pathway to upregulate TIMP-3 expression in SMCs. The increased TIMP-3 reduces MMP2/9 activity, thus preventing collagen and elastin from degrading within the plaques. Mechanistically, we found that nuclear STAT6 recruits the coactivator CBP/p300 to bind the TIMP-3 promoter. The interaction with CBP/p300 increases acetylation of STAT6 and TIMP-3 promoter histones, which initiates TIMP-3 transcription. Our studies support clinical observations and demonstrate that IL-16 has a protective role in atherosclerotic plaque stabilization. ^17^

Atherosclerotic plaque rupture and subsequent thrombus generation triggering ischemic heart disease, such as myocardial infarction, are the main causes of mortality. Improving plaque composition and increasing plaque stability is an effective strategy for reducing acute coronary syndrome. The features of vulnerable plaques include a high content of macrophages and lipids and less SMCs and collagen. Reducing lipid deposits and inhibiting inflammatory cell infiltration in the subendothelial space can demonstrably inhibit the development of atherosclerosis and decrease plaque vulnerability.^19^ On the other hand, a high content of SMCs and collagen forming a thick fibrous cap also contributes to plaque stability. Increasing the synthesis of collagen by various compounds, such as thiazolidinediones and dapagliflozin, or inhibiting the degradation of collagen by matrix metalloproteinases (MMPs) leads to a more stable phenotype of arteriosclerotic plaques.^20-22^ In this study, we showed that IL-16 upregulated TIMP-3 and reduced MMP activities, which increased collagen content as well as plaque stability. Therefore, IL-16 was demonstrated here to be a novel factor involved in the modulation of the extracellular matrix and plaque phenotype, which is the most important finding of this study.

Due to its unique ability to bind to ECM and inhibit a broad range of inhibitory substrates, including MMPs, ADAMs, and ADAMTSs, TIMP-3 is recognized as a crucial regulator of ECM remodeling among the four TIMPs.^1,2^ TIMP-3 transcription is strictly regulated to prevent excessive ECM degradation and protect tissue function. The TIMP-3 promoter region reportedly contains putative consensus binding sequences of multiple transcription factors, such as Sp1, Smad2/3/4, AP-1 NF-κB, c-Myc, and p53.^23,24^ TIMP-3 expression responds to transforming growth factor-β1 (TGF-β1), which stimulates TIMP-3 expression by promoting the direct binding of Smad2/3/4 to the conserved sites of the TIMP-3 promoter.^25^ In addition to Smads, TGF-β1 and bovine lactoferricin can also increase ERK1/2-mediated Sp-1 activation to upregulate TIMP-3 in chondrocytes.^26,27^ Here, we provided evidence that the cytokine IL-16 can stimulate TIMP-3 expression, and the upregulation requires binding of STAT6 to its promoter in VSMCs. Moreover, we further found that the histone acetyltransferase CBP/P300 acts as a transcriptional coactivator of STAT6 for transcriptional activation of the TIMP-3 genes by increasing histone acetylation in gene loci. Recently, downregulation of histone deacetylase 9 (HDAC9) was reported to increase TIMP-3 promoter histone acetylation and TIMP-3 expression levels in trophoblast cells, suggesting an impact of acetylation on TIMP-3 transcription.^28^ Here, we extend the previous findings and reveal a direct role of CBP/p300 in promoting TIMP-3 transcription.

In addition to the acetylation modification that is involved in TIMP-3 transcription regulation, as discussed above, TIMP-3 is reportedly also controlled by microRNAs such as microRNA-181b, microRNA-222, and microRNA-21.^29-31^ Moreover, the cytokines IL-1β, IL-6, IL-17 and IL-23 modulate viral infection, pulmonary hypertension, the inflammatory response, and collagen production in scleroderma fibroblasts by regulating various microRNAs.^32-35^ Therefore, we could not rule out the possibility of the involvement of microRNAs in IL-16-mediated TIMP-3 expression.

IL-16 is induced by antigen and mitogen stimulation or virus infection in T lymphocytes and acts as a lymphocyte chemoattractant factor to increase immune cell migration and recruitment into inflammatory lesions, promoting the synthesis of proinflammatory cytokines, including IL-1 and IL-6, and tumor necrosis. Therefore, in this respect, IL-16 seems to enhance atherosclerotic development. Overexpression of IL-16 augmented cardiac inflammation, leading to increased cardiac fibrosis, while inhibition of IL-16 by administering an IL-16 neutralizing antibody attenuated doxorubicin-induced cardiac injury^14,16^. However, we observed that IL-16 conferred beneficial effects on vascular lesions in this study, which is consistent with a clinical investigation showing that a high level of IL-16 is positively correlated with a reduced risk for cardiovascular events,^17^ suggesting the existence of an alternative mechanism underlying IL-16 function. Here, we showed that IL-16 increases TIMP-3 production in VSMCs, leading to plaque stabilization. Moreover, in addition to its role in maintaining ECM homeostasis, TIMP-3 has been demonstrated to reduce atherogenesis by inhibiting macrophage activation,^31, 36-38^ which arises from at least the following two aspects. First, TIMP-3 has been demonstrated to inhibit TNF-α convertase (TACE)/ADAM17 and TNF-α shedding from the membrane-bound form, which reduces the soluble TNF-α-induced inflammatory response, including activation of NF-kB and IL-6 production in macrophages.^39,40^ TIMP-3-deficient mice reportedly exhibited increased inflammatory markers and M1 macrophage polarization in the aorta and showed more severe atherosclerotic lesions.^36^ Second, TIMP-3 can directly inhibit macrophage invasion and reduce proliferation and apoptosis, which destabilizes atherosclerotic plaques.^37,38^ Therefore, the impacts of macrophage activation induced by IL-16 are complex and should be taken into account when we fully elucidate the role of IL-16 in cardiovascular disease. Furthermore, based on the discussion above, it is interesting to explore whether IL16 has opposite effects at specific stages of atherosclerosis development. For example, increased circulating IL-16 triggers macrophage infiltration and accumulation under the intima, which promotes atherosclerosis development at the early stage, while in the later stage, the production of TIMP-3 in SMCs stimulated by IL-16 inhibits macrophage activation and plaque phenotypic remodeling at lesion sites.

Maintenance of arterial ECM integrity is primarily important for vascular function. Rearrangement of arterial ECM components, particularly collagen and elastin, induces vascular remodeling and is closely related to vascular diseases such as abdominal aortic aneurysm (AAA) and hypertension. Evidence shows that the deletion of TIMP-3 promotes AAA formation and hypertension and decreases the survival rate in an angiotensin II-infused murine model.^3,4,31,41^ Here, we demonstrated the involvement of IL16 in ECM remodeling by upregulating TIMP-3. Therefore, it is reasonable to assume that in addition to atherosclerosis, IL-16 might also have a protective role in AAA and hypertension.

The aortic MMP2/9 gelatinolytic activities were increased in IL-16-defective mice in this study, although their expression was not significantly altered (data not shown). In isolated VSMCs, we found that the expression and enzymatic activities of MMP2/9 were increased in response to IL-16 stimulation; the expression of proliferating cell nuclear antigen (PCNA) was not significantly changed (data not shown). As a result, VSMC migration was promoted by IL-16 with no effects on proliferation, which is in line with previous reports (data not shown).^42^ VSMC migration and invasion from the media into the intima contribute to thick fibrous cap formation and plaque stabilization in advanced atherosclerosis. This finding may explain our observation that a reduced amount of VSMCs was detected within the plaques in IL16-/-ApoE-/- mice.

Although CD4 is a well-recognized receptor for IL-16, other molecules, such as CD9, have been reported to serve as alternative receptors for IL-16.^43,44^ Therefore, whether the IL-16-directed upregulation of STAT6/TIMP-3 revealed in this study is also mediated by other receptors remains to be further investigated. In addition, we note that cellular signaling molecules such as c-Jun N-terminal kinase (JNK), stress-activated protein kinase (SAPK) and p38 are involved in the IL-16-induced pathway.^45,46^ We also detected an increase in the tyrosine-phosphorylated Src family member Lyn, but not Fyn, in IL-16-induced VSMCs (data not shown). In this study, we highlight nuclear events, especially TIMP-3 transcriptional regulation. Therefore, it is of interest to further investigate whether the effects of IL-16 on TIMP-3 transcription are dependent on such cellular kinases and clarify the detailed mechanism of upstream signal transduction.

In summary, IL-16 prevents extracellular matrix degradation and enhances plaque stability by upregulating TIMP-3 in smooth muscle cells. Activation of the downstream signaling pathway CD4/JAK2/STAT6 and cooperation with CBP/p300 are required for IL-16-induced TIMP-3 expression. Our findings suggest that IL-16 is a novel factor in vascular remodeling and atherosclerotic plaque phenotype modulation, which furthers our understanding of atherosclerotic development in later stages and provides us with clues in finding an effective strategy to reduce clinical incidences caused by plaque rupture.

## Non-standard Abbreviations and Acronyms

IL-16: interleukin 16
TIMP-3: tissue inhibitors of metalloproteinase 3
ECM: extracellular matrix
MMPs: matrix metalloproteinases
HFD: high-fat diet
CBP: cAMP-response element binding protein
BCA: brachiocephalic artery
ColIα1: collagen I α1

## Supplemental Material

Supplemental Methods

Tables S1–S5

Figure S1–S9

